# Characterising sex differences of autosomal DNA methylation in whole blood using the Illumina EPIC array

**DOI:** 10.1101/2021.09.02.458717

**Authors:** Olivia A Grant, Yucheng Wang, Meena Kumari, Nicolae Radu Zabet, Leonard Schalkwyk

## Abstract

Sex differences are known to play a role in disease etiology, progression and outcome. Previous studies have revealed autosomal epigenetic differences between males and females in some tissues, including differences in DNA methylation patterns. Here, we report for the first time an analysis of autosomal sex differences in DNAme using the Illumina EPIC array in human whole blood (n=1171). We identified 554 sex-associated differentially methylated CpG sites (saDMPs) with the majority found to be hypermethylated in females (70%). These saDMP’s are enriched in CpG islands and CpG shores and located preferentially at 5’UTRs, 3’UTRs and enhancers. Additionally, we identified 311 significant sex associated differentially methylated regions (saDMRs). Transcription factor binding site enrichment revealed enrichment of transcription factors related to critical developmental processes and sex determination such as SRY and SOX9. Our study reports a reliable catalogue of sex associated CpG sites and elucidates several characteristics of these sites.

## INTRODUCTION

Sex is an important covariate in all epigenetic research due to its role in the incidence, progression and outcome of many phenotypic characteristics and human diseases (1, 2). There is an increasing interest as to which role epigenetic modifications (such as DNA methylation) may play in the underpinnings for relationships between environmental exposures and disease onset. In addition, sex has previously been shown to have a strong influence on DNA methylation variation (3–7). However, the idea that DNA methylation variation between males and females may underlie the sex biases observed in diseases has not been well documented thus far.

Sex differences in disease prevalence are sometimes explained at the molecular level and rooted in genetic differences between males and females. Differences in sex chromosome complement have independently been shown to direct differences in gene expression and chromatin organization (8–11). Furthermore, these differences in sex chromosome complement are sufficient to explain sex bias seen in some diseases. X chromosome number has previously been shown to impact immune cell population and occasionally therefore the development of diseases such as autoimmunity (12, 13).

Previous research has also revealed sex differences in gene expression of autosomal genes as well as sex chromosome linked genes (14). It is worth noting that most of the differences in gene expression on the autosomes are small differences (15). However small expression differences may still be associated with great effects on phenotypic characteristics and disease incidence and onset. Others also identified sex differences in chromatin accessibility and histone modifications, thus suggesting that different epigenetic factors contribute to gene expression sex biases seen in some diseases (16).

Sex specific gene expression and levels of sex hormones may be mediated by epigenetic mechanisms, including DNA methylation. Several genome wide association methylome studies (or Epigenome Wide Association Studies - EWAS) have highlighted differences in DNA methylation patterns linked to sex differences in genes on the autosomes (17–20). Previous studies have reported sites showing varying methylation due to sex differences in various tissues such as saliva, placenta, brain, pancreatic islets and whole blood (17, 19, 21–27). Early studies in this area did not account for cross hybridizing probes which map to both autosomal and sex chromosome loci (28, 29). A meta-analysis of 76 published studies with correction for cross hybridizing probes only validated 184 sex associated sites (30). Furthermore, cellular heterogeneity is also important for identification of sex associated sites as several studies have previously demonstrated that cell type proportions may occasionally vary by sex and that could result in identification of cell type associated type differences rather than sex associated differences (24).

Due to X chromosome inactivation in females, large differences in methylation levels of X chromosomes can be observed between males and females (31). Recent research suggests that normalising methylation data with the sex chromosomes introduces a large technical bias to many autosomal CpGs. This technical bias has been reported to result in many autosomal CpG sites being falsely associated with sex and extra steps need to be taken to ensure minimal technical bias is introduced to the autosomal CpGs.

Here, we use the EPIC BeadChip to assess autosomal sex differences in DNA methylation levels from whole blood at individual sites and genomic regions. All individuals involved in this study were part of Understanding Society: The UK Household longitudinal study (32). Additionally, we account for cell type composition, cross hybridization and adequately handle the technical bias introduced by sex chromosomes. To our knowledge, this is the first study using the Illumina EPIC BeadChip (allowing for interrogation of ~850,000 sites across the genome) to investigate autosomal sex differences in DNA methylation.

## MATERIAL AND METHODS

### Participants

Whole blood Illumina Infinium MethylationEPIC BeadChip DNAme data was collected from 1175 participants involved in Understanding Society: The UK Household Longitudinal Study (33). In wave 3 of the study (2011-12) blood samples were collected from a portion of the study participants. Individuals were considered eligible to give a blood sample if they were over the age of 16 and met the other requirements detailed in the UK Household Longitudinal Study Biomarker user guide. Our study population was restricted to participants of white ethnicity. A full description of the dataset and data processing has been described by (34).

### DNA methylation data

Samples of whole blood DNA from 1175 participants were obtained following the protocol described in (34). Raw signal intensities were processed using the R package bigmelon (35) and wateRmelon (36)from idat files. Prior to normalisation of the data, outlier samples were identified using principal component analysis and subsequently removed from the data set. The reported age of each sample was compared to predicted age using the epigenetic age method implemented by *agep* in the R package bigmelon (35) .Further, the reported sex of the samples was checked using a DNA methylation-based sex classifier (31)which predicts sex based on the methylation difference of X and Y chromosomes. 4 samples were subsequently removed after this step as reported and predicted sex did not match. The data was then normalised via the *interpolatedXY* adjusted *dasen* method using the *adjustedDasen* function in the R package watermelon (37) which first normalises autosomal CpGs by *dasen* and then the corrected methylation values of sex chromosome linked CpGs are estimated as the weighted average of their nearest neighbours on autosomes. Following normalisation of the data, SNP probes, cross hybridizing probes (29) and X or Y linked probes were removed from the data set. The final data set consisted of 1171 samples and 747,361 DNA methylation sites.

To ensure that whole blood cell composition did not differ significantly by sex and would not introduce bias to our results, the relative proportions of Granulocytes, mononuclear, natural killer, CD4T, CD8T and B cells were estimated for all samples using the *estimateCellCounts* function implemented in bigmelon (35). Furthermore, to assess whether the sex differences we observed were age independent, we performed a Mann-Whitney U test between the age distribution of males and females. Our results confirmed that there is no statistical difference in age between our male and female samples (p value 0.07; median values of 60 and 58, respectively).

### Identifying sex associated autosomal differential methylation

Sex associated autosomal differentially methylated positions (saDMPs) were identified by performing linear modelling using the ChAMP package in R (38) using sex and Beta values. Correction for multiple testing was performed with the Benajamini-Hochberg false discovery rate method (FDR values). A probe was considered significantly differentially methylated if the difference in Beta values between males and females was greater than 0.05 in either direction and the FDR value was smaller than 0.05. We further characterized differentially methylated regions (DMRs) by applying the *DMRcate* function from the R package ChAMP (38) to detect DMRs between males and females on the autosomes. A DMR was considered to be significantly associated with sex (saDMR) if the FDR value was smaller than 0.05 and consisted of at least 2 CpG sites with a difference in beta values between males and females greater than 0.05.

### Genomic annotation of CpG sites

We annotated the autosomal CpG’s using the manufacturer supplied annotation data (MethylationEPIC_v-1-0_B2 manifest file). Annotation was completed in the R package Minfi (39). Several categories were used as annotations in relation to CpG islands and divided into the following categories: CGIs, CGI shores (S and N), CGI shelfs (S and N) and open sea regions. Further, we also annotated the autosomal CpGs to several genomic features, including exons, introns, 5’ UTR, 3’UTR, enhancers, promoters and transposable elements (TEs) using data from UCSC table browser (https://genome.ucsc.edu/cgi-bin/hgTables).

### Gene ontology analyses

GO analyses were conducted using the *gometh* function in the missMethyl package (40) which tests gene ontology enrichment for significant CpGs while accounting for the differing number of probes per gene present on the EPIC array. For GO ontology analyses of enriched TFBS we used *enrichGO* from the clusterProfiler package in R (41), which performs FDR adjustment.

### Enrichment of saDMPs in transcription factor binding motifs and integration with gene expression

The enrichment analysis of known motifs in sex associated DMPs was performed using the R package PWMEnrich (42) using the MotifDb collection of TF motifs (43). Specifically, the DNA sequences within a 100 bp range from the saDMP which were hypermethylated in females were extracted from the genome and compared to the saDMPs which were hypermethylated in males as the background to reveal unique enriched motifs (adjusted p-value < 0.05). RNA-seq data for 20 healthy donors (10 males and 10 females) from publicly available data from GEO (GSE120312) was used in our analysis. In particular, we used the pre-processed count matrices with DESeq2 (44) to calculate differentially expressed genes between males and females with an adjusted p-value of 0.05 and log_2_ fold change of 1.

### Overlap of saDMP’s with chromatin loops

We examined whether any of the sex associated DMPs made 3D contacts with distal genes using Hi-C data available from the Gene Expression Omnibus (GEO) under accession number (GSE124974) for white blood cells and neutrophils. Hi-C library preparation was performed using the Arima-HiC kit and pre-processing of the data was performed using Juicer command line tools (45). Reads were aligned to the human (hg38) genome using BWA-mem (46) and then pre-processed using the Juicer pre-processing pipeline. We called chromatin loops using the HICCUPS tool from Juicer using a 10 Kb resolution. We then constructed GenomicInteractions objects to annotate saDMPs to loop anchors using the *findOverlaps* function from the GenomicRanges package using a *maxgap* of 10000. Following this, we then annotated the corresponding anchor to the relevant gene ID. These steps then allowed us to perform network analysis in Cytoscape (47) and GO and KEGG analyses in clusterProfiler (41).

### Protein-protein network visualisation and hub gene identification

We searched all of the genes annotated to our saDMPs using the Search Tool for the Retrieval of Interacting Genes (STRING) (https://string-db.org) database to generate our networks. We extracted protein-protein interactions with a combined score of >0.4. Then, we extracted top 50 genes using degree of connectivity methods in the plugin tool CytoHubba. The same analysis was repeated for the enriched transcription factor motifs found at saDMPs.

## RESULTS

### Females show higher methylation at a subset of autosomal loci

Analysis of DNA methylation (DNAme) differences between males and females on the autosomes was performed using linear regression for the IlluminaEPIC BeadChip for 1171 individuals (682 females and 489 males). After data processing and cleaning, n=747,302 CpGs were analysed (see *Material and Methods*). Sites which are known SNP probes, cross hybridizing or X/Y linked probes were excluded. After adjusting for multiple testing using the Benjamini Hochberg FDR method (FDR p < 0.05) we identified 14,653 CpGs associated with sex. Of those CpGs, 68% (9960 CpGs) were more highly methylated in females and the remaining 32% (4693 CpGs) were more methylated in males. Gene ontology analyses showed several enriched terms for these 14,653 CpGs (Table 1) which included many terms related to development such as cellular developmental processes, system development, anatomical structure development and multicellular organism development. Furthermore, other enriched terms included plasma membrane, ion transport and more (see Table 1).

**Table 1.** Enriched GO terms among the 14,653 CpGs identified to be significantly associated with sex. N^1^ indicates the number of genes in the GO term. DE^2^ refers to the number of genes annotated to the sex associated DMPs which are differentially methylated. P.DE^3^ indicates the P value for over representation of the GO term in this data set. FDR^4^ indicates the false discovery rate (using the Benjamini and Hochberg method).

| GO term | Term | Ont | N<sup>1</sup> | DE<sup>2</sup> | P.DE<sup>3</sup> | FDR<sup>4</sup> |
| --- | --- | --- | --- | --- | --- | --- |
| GO:0048856 | anatomical structure development | BP | 5493 | 1820 | 1.27E-12 | 2.53E-08 |
| GO:0044459 | plasma membrane part | CC | 2610 | 934 | 2.24E-12 | 2.53E-08 |
| GO:0071944 | cell periphery | CC | 5023 | 1616 | 4.32E-12 | 3.26E-08 |
| GO:0005886 | plasma membrane | CC | 4919 | 1580 | 6.86E-12 | 3.88E-08 |
| GO:0007275 | multicellular organism development | BP | 5048 | 1677 | 1.10E-11 | 4.19E-08 |
| GO:0048468 | cell development | BP | 1999 | 768 | 1.14E-11 | 4.19E-08 |
| GO:0032501 | multicellular organismal process | BP | 7000 | 2160 | 1.30E-11 | 4.19E-08 |
| GO:0032502 | developmental process | BP | 5848 | 1902 | 1.49E-10 | 4.21E-07 |
| GO:0009653 | anatomical structure morphogenesis | BP | 2493 | 910 | 3.23E-10 | 8.11E-07 |
| GO:0048731 | system development | BP | 4537 | 1506 | 1.22E-09 | 2.64E-06 |
| GO:0030154 | cell differentiation | BP | 3882 | 1294 | 1.28E-09 | 2.64E-06 |
| GO:0043269 | regulation of ion transport | BP | 628 | 255 | 2.98E-09 | 5.61E-06 |
| GO:0006811 | ion transport | BP | 1543 | 550 | 6.78E-09 | 1.18E-05 |
| GO:0048869 | cellular developmental process | BP | 4054 | 1339 | 9.49E-09 | 1.53E-05 |
| GO:0009887 | animal organ morphogenesis | BP | 957 | 384 | 1.23E-08 | 1.86E-05 |
| GO:0016020 | membrane | CC | 8539 | 2569 | 1.74E-08 | 2.46E-05 |
| GO:0098916 | anterograde trans-synaptic signaling | BP | 663 | 276 | 2.50E-08 | 3.14E-05 |
| GO:0007268 | chemical synaptic transmission | BP | 663 | 276 | 2.50E-08 | 3.14E-05 |
| GO:0044425 | membrane part | CC | 6238 | 1866 | 4.20E-08 | 5.00E-05 |
| GO:0099537 | trans-synaptic signaling | BP | 670 | 277 | 5.96E-08 | 6.73E-05 |

The lambda value of the Q-Q plot is 1.99 (Figure S1A) indicating slight inflation of test statistics and, in order to ensure we detect true sex differences, we selected CpGs that displayed large differences in methylation. Thus, we further filtered our list of 14,653 CpGs by only considering those probes that displayed a ΔBeta value (absolute difference between average Beta values in male and female samples) greater than 0.05. A total of 554 CpGs met this criterion (called sex associated DMPs or saDMPs) and were distributed across all autosomes (Figure 1A). 70% of the saDMPs were hypermethylated in females (389 CpGs) and 30% of which were hypermethylated in males (166 CpGs) (Figure 1B) (See *Additional File 1* for the full list). Moreover, since whole blood is a bulk tissue, we calculated the estimated cell type proportions for whole blood between our male and female samples to assess whether any differences in cell type proportions would potentially be reflected in our results resulting in false positives. Using Wilcoxon test, we found no significant difference in the proportions of Granulocytes and CD4T cells between males and females, but we did find statistically significant differences in proportions of CD8T, Natural killer, B cells and monocytes (Supplementary figure 1B). We therefore performed our analysis with and without accounting for cell type proportions to see how this would affect our results. We found that we still retrieved the same list of saDMPs with and without accounting for cell type proportions allowing us to conclude that differences in cell type proportions did not affect our analysis.

**Figure 1:**
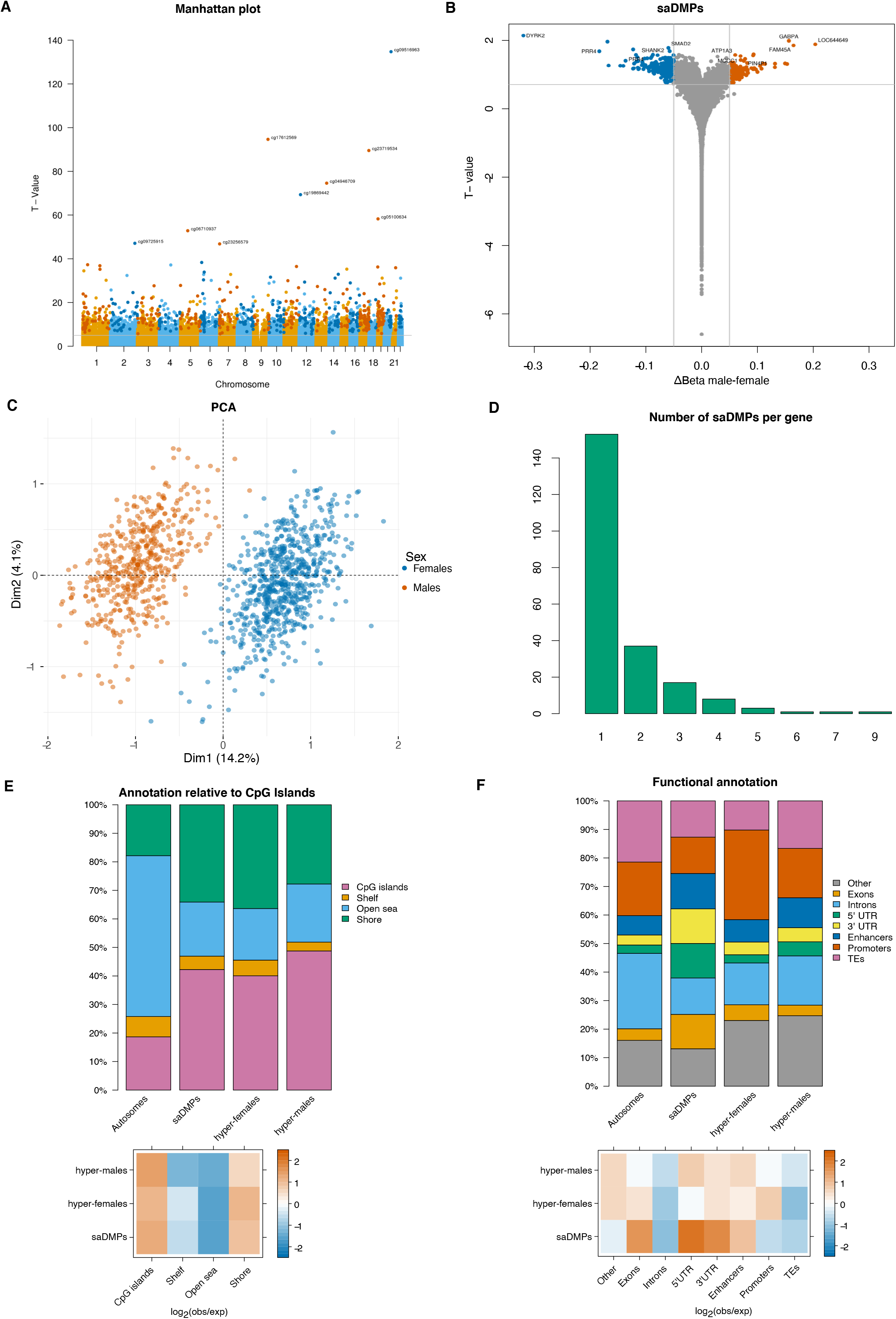
Location and characterisation of saDMPs. (A) Manhattan plot for EWAS analysis of sex. CpG sites which met a threshold of FDR < 0.05 and had an average beta change of > 0.05 were considered significant and are represented by darker colours. (B) Volcano plot for saDMPs. CpGs which are not significant are represented in grey, saDMPs hypermethylated in males are in orange and saDMPs hypermethylated in females in blue. (C) Principal component analysis of beta values at the significant saDMPs. Male samples are indicated in orange while female samples are indicated in blue. (D) Number of saDMPs harboured by individual genes. (E) Top panel shows the annotation of all saDMPs (n=544), saDMPs hypermethylated in females (n=382) and saDMPs hypermethylated in males (n=162) relative to CpG island regions compared to the autosomal background. Bottom panel shows the log_2_ (obs/exp) annotations based on the autosomal background of the different annotations. (F) Top panel shows the overlap of all saDMPs (n=544), saDMPs hypermethylated in females (n=382) and saDMPs hypermethylated in males (n=162) with genomic features compared to the autosomal background. Bottom panel shows the log_2_ (obs/exp) annotations based on the autosomal background of the different annotations.

Since we had such stringent parameters to define what we considered a significantly associated saDMP for males and females. We performed principal component analysis (PCA) to see how male and female beta values clustered in PC space and to evaluate the effect of DNAme at the saDMPs. As shown in Figure 1C, male and female samples formed clear clusters based on the beta values of the significant sex associated DMPs (554 CpGs). PC1 explained 14.2% of the variance and PC2 explained 4.1% of the variance. Based on Figure 1C we can conclude that these saDMPs are sufficient to contribute to the clear separation of male and female samples in PC space.

### Characterisation of sex associated DMPs

The saDMPs were found in 223 unique genes with 68 of these genes harbouring several saDMPs with an average of 2.7 saDMPs per gene (Figure 1D). CRISP2, a gene known to be involved in sperm function and male fertility (Lim et al.2019), harboured the largest number of saDMPs, 9. We performed GO and KEGG analyses, but did not identify any significantly enriched biological processes or pathways for these genes.

To help us try to gain more insight into the functional role of these saDMPs, we characterised their genomic location and further compared this with the autosomal background. We found saDMPs are preferentially located CpG islands and CpG shores and depleted in open sea regions compared to the autosomal background (Figure 1E). Moreover, saDMPs hypermethylated in females are enriched at promoters and exons, with saDMPs hypermethylated in males being enriched at 5’UTR, potentially acting as alternative promoters (Figure 1F). Interestingly, we observed that all saDMPs display enrichment at enhancers, which, together with their presence at promoters, indicates they could play a role in gene regulation. Lastly, we also note that all saDMPs were depleted at transposable elements and introns compared to the autosomal background.

Enrichment of saDMPs at enhancers suggests that some of the saDMPs could potentially regulate distal genes (48, 49). We further annotated the saDMPs to genes by assessing their contacts with promoters, and thus, distal genes. Following this, we further annotated the saDMPs to 46 additional genes, 35 of them being annotated to saDMPs hypermethylated in females and 11 to saDMPs hypermethylated in males (see Figure S2A-B).

To evaluate the interactions between the proximal (located in genes) and distal (those in 3D proximity) saDMPs, we collated these and produced protein-protein interaction networks to visualize the networks of these genes (see Figure S2C-D). Of the 11 genes linked to saDMPs hypermethylated in males, we found three histones (HIST1H3A, HIST1H4A and HIST1H4B), which are known to interact with CDYL (Figure S2C), a gene that we also found to harbour an saDMP hypermethylated in males. Chromodomain Y-like protein (CDYL) is a chromatin reader binding to heterochromatin (H3K9me3, H3K27me2 and H3K27me3) that is crucial for spermatogenesis, male fertility and X chromosome inactivation (50). From the list of genes linked to saDMPs hypermethylated in females, KDM2A regulates circadian gene expression by repressing activity of CLOCK-ARNTL which has previously been shown to exhibit sex dimorphism (51). In addition, ODF2L; outer dense fiber of sperm tails 2 like is also linked to saDMPs hypermethylated in females and has previously be shown to interact with PRSS23, which is involved in ovulation (52).

We then performed functional enrichment of genes annotated to those saDMPs to identify enriched biological processes or terms. Enrichment of the genes annotated to those saDMPs hypermethylated in females revealed several processes (129 terms), including adaptive immune system, estrogen receptor binding and androgen receptor binding (FDR < 0.05) (*Additional File 5*). Furthermore, enrichment of the genes annotated to those saDMPs hypermethylated in males displayed enrichment of terms such as dosage compensation by inactivation of X chromosome, hormone receptor binding and gene / chromatin silencing (FDR <0.05) (*Additional File 6*).

### Enrichment of saDMPs in transcription factor binding sites

To identify common features among the sex associated DMPs, we performed transcription factor (TF) binding site and gene ontology analyses. First, we evaluated whether the saDMPs were enriched in motifs for TFs (100 bp window). For the saDMPs hypermethylated in females, we found 329 enriched TFs (p.value < 0.05) (Figure 2A and *Additional File 3*) with strongest evidence for TROVE2, XRCC1 and NFIL3. TROVE2/R060 is an RNA binding protein known to be involved in stimulating of androgen receptor regulated genes. We also found SOX9 and SRY TFs to be enriched in DMPs hypermethylated in females, which are genes known to be involved in male sex determination (Figure 2A) (53). For saDMPs hypermethylated in males, we identified 64 enriched TFs, including CEPBG, AFF4 and ELK1. CEPBG is a transcription factor which has previously been reported to stimulate adipocyte differentiation in an ESR1/CEBPA mediated pathway (54). Furthermore, GABPA and ELK1 have previously been shown to be associated with sex associated differentially methylated regions in the mouse liver (55).

**Figure 2:**
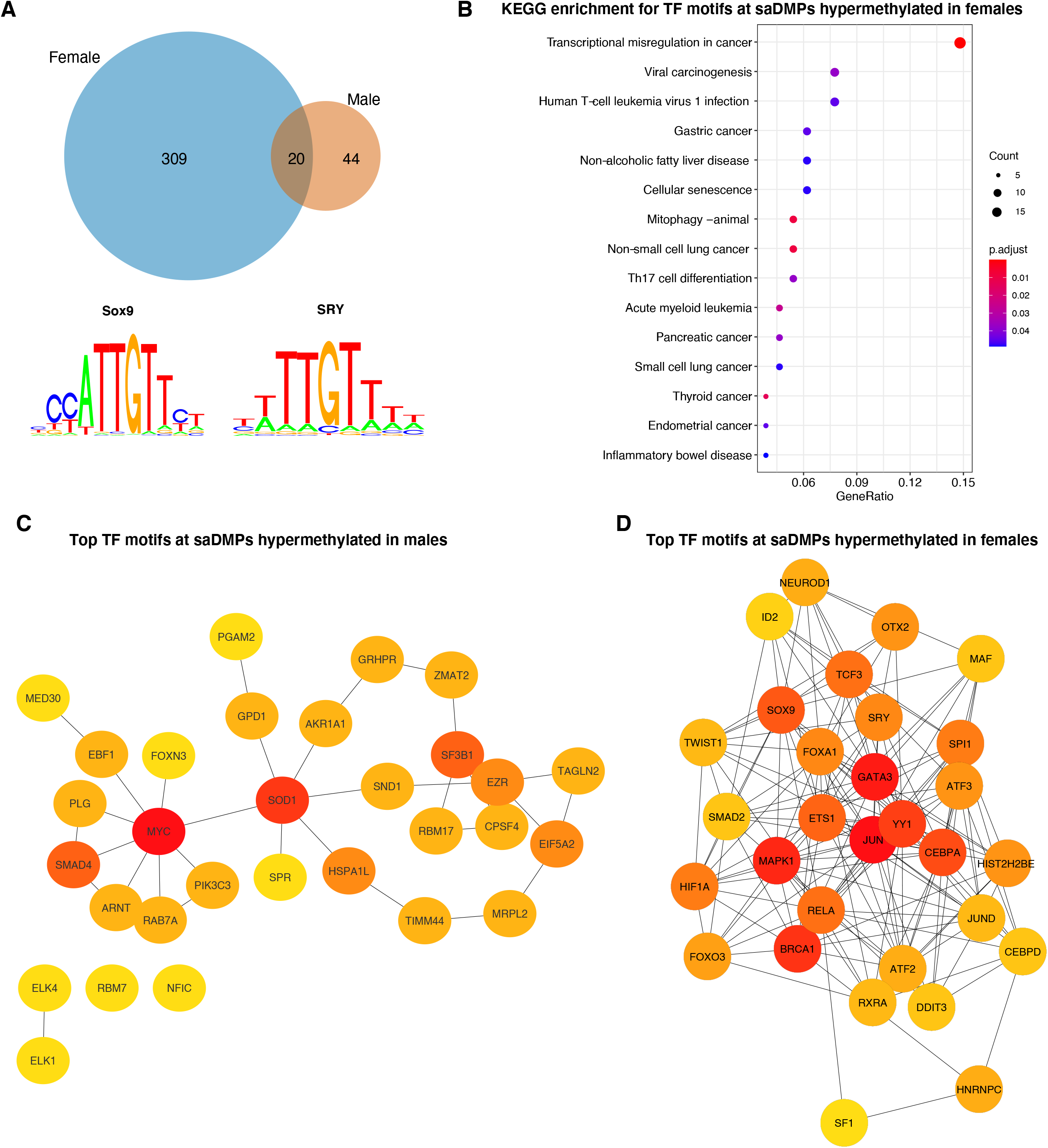
Transcription factor motif enrichment analysis. (A) Overlap of enriched TF motifs for saDMPs hypermethylated in females (blue) and saDMPs hypermethylated in males (orange). saDMPs found to be hypermethylated in females were enriched in TF binding motifs including SOX9 and SRY. (B) KEGG analyses for the significantly enriched TF motifs at saDMPs hypermethylated in females. (C-D) Subnetworks of the top 30 enriched TF motifs at saDMPs hypermethylated in males (C) and females (D). Node colour represents the degree of connectivity. The scale from red to yellow represents the top 30 enriched TF motif rank from 1-30, with red indicating highest degree and yellow indicating lowest degree.

To analyse whether the TF motifs were enriched for annotation to biological processes or pathways, we performed pathway analyses using the GO and KEGG databases. We identified 15 enriched KEGG pathways for the TFBS enriched at saDMPs hypermethylated in females, spanning a wide range of processes such as the transcriptional regulation in cancer, several specific cancer pathways, viral carcinogensis and more (Figure 2B). In addition, we also found 39 enriched GO terms ranging from transcription factor activity, E-box binding, transcription coactivator activity and interestingly, bHLH transcription factor binding (Figure S3B). Nevertheless, we found no enriched KEGG terms for the TFs enriched at saDMPs hypermethylated in males, likely due to the small number of enriched TFs. However, we identified 9 enriched GO terms such as NAD, NADP binding and oxidoreductase activity (Figure 3SA).

**Figure 3.**
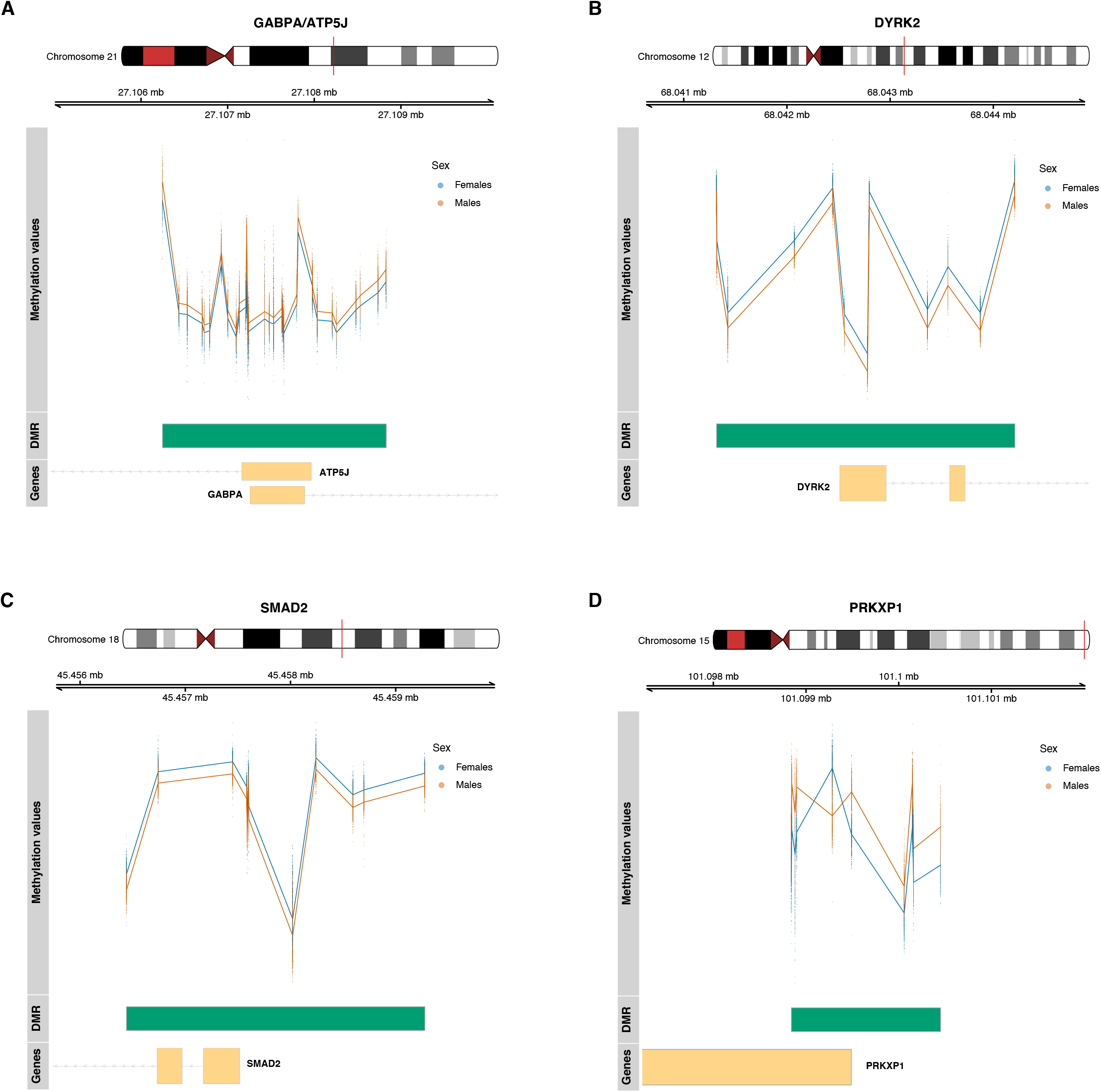
Plots of sex associated differentially methylated regions (saDMR). We plotted regions: (A) GABPA and ATP5J, (B) DYRK2, (C) SMAD2 and (D) PRKXP1. Yellow boxes represent appropriately labelled genes, green boxes represent the genomic region which the differentially methylated region spans. The scatterplots represent the methylation values for males (orange) and females (blue) at CpG sites located within the differentially methylated region.

One possibility is that transcription factor motifs for genes encoded on the sex chromosomes may act as hubs in the enriched TF motif network. To assess this, we produced protein-protein interaction networks to visualize the networks of these TFs (Figure S4). Although we identified some enriched motifs for several TFs encoded on the X chromosomes in the saDMPs hypermethylated in males such as ELK1, TGIF2LX and TCEAL6 (Figure S4A), we observed that they were not central hub nodes in the network. Nevertheless, we did identify several central TF motifs encoded on the sex chromosomes for the saDMPs hypermethylated in females (Figure S4B). These included 11 TFs encoded on the X chromosome and 2 on the Y chromosome including SRY and KDM5D.

We further utilised cytohubba package to robustly identify if these TFs were in fact hub genes in the network. This revealed that ELK1 did in fact act as a hub gene in the TF network, however the other 49 genes were encoded on the autosomes (Figure 2C). Interestingly, for the TF motifs enriched at saDMPs hypermethylated in females, we found SRY (coded for on the Y chromosome) and SOX9, BRCA1 and other autosomal genes (Figure 2D). BRCA1 is a tumour suppressor gene which is heavily implicated in breast cancer and its promoter hypermethylation in peripheral blood has previously been shown to be linked to decreased risk of breast cancer (56).

### Relationship with gene expression

The 554 saDMPs were then further explored in association with the expression levels of their annotated genes using publicly available data for whole blood poly(A)+. We did not identify any sex biased gene expression patterns corresponding to differences in DNAme levels at these genes (Figure S5). Some of the genes annotated to our saDMPs met the P value threshold after correcting for multiple testing using the Benjamini and Hochberg method (adjusted pvalue<0.05), however did not meet the log fold change threshold (log_2_FC>1) such as TARP, KCNT1 and CEBPB (Figure S5A). However, these genes have low expression in blood and therefore, this analysis may be more relevant for other tissues such as brain or skeletal muscle. The majority of the differentially expressed genes are located on the sex chromosomes, but we also did observe differential expression between males and females for several autosomal genes (Figure S5B-S5C). These results indicate that the differences in DNA methylation observed between males and females do not lead to any large difference in gene expression, but only some small ones.

### Sex associated differentially methylated regions

Given that several genes harboured numerous saDMPs, we postulated whether some of the saDMPs were part of larger differentially methylated regions associated with sex. We therefore searched for differentially methylated regions associated with sex. Following adjustment for multiple testing (FDR), we identified a large list of 14,386 sex associated differentially methylated regions. We therefore considered a saDMRs significant if it harboured at least 2 CpGs, had an FDR value smaller than 0.05 and had a maximum absolute beta fold change value within the region greater than 0.05 in either direction. Following filtering of the list of saDMRs, we identified 311 significant sex associated DMRs on the autosomes between males and females located in 226 unique genes (*Additional File 2*). The number of CpGs within the DMRs ranged from 3 to 58 and had an average width of 1793 base pairs (bp) ranging from 63bp to 5638 bp.

Figure 3 shows the methylation values for males and females at 4 of the most significant saDMRs: *(i)* the strongest saDMR (31 CpGs) overlaps ATP5J and GABPA genes on chromosome 21 (Figure 3A), *(ii)* the second most significant saDMR (12 CpGs) overlaps DYRK2 gene on chromosome 12 (Figure 3B), *(iii)* the third strongest DMR (12 CpGs) overlaps SMAD2 gene on chromosome 18 (Figure 3C) and *(iv)* the fourth strongest saDMR (10 CpGs) overlaps PRKXP1 pseudogene on chromosome 15 (Figure 3D). DYRK2 and SMAD2 are hypermethylated in females, while ATP5J, GABPA and PRKXP1 are hypermethylated in males. Note that PRKXP1 saDMR, although hypermethylated in males, contains a single CpG which is hypermethylated in females.

A saDMR harbouring 58 CpGs overlapped the promoter region of a gene called SCAND3. The top hits in the saDMR list overlapped promoter regions of the following genes; ATP5J, GABPA, DYRK2, SMAD2, CRISP2, AND PRKXP1. ATP5J and GABPA are genes (hypermethylated in males) which have previously been reported to be implicated in early onset of Alzheimers disease (57), a disease known to affect females more than males. Furthermore, ATP5J is a gene known to be a target gene of oestrogen, previously shown to serve an inhibitory role in the sex differences in hepatocellular carcinoma (58). DRYK2, SMAD2 and CRIPS2 have previously been shown to exhibit functions which are sex specific (21). CpGs harboured by the gene ZPBP2 are hypermethylated in females and previous work has implicated this gene in sex specific effects in asthma and, consistent with our findings, also showed that methylation at this gene is lower in males than females. PRKXP1 is located on chromosome 15 and CpGs in this region have previously been associated with Crohns disease and intestinal inflammation, a disease which has previously been reported to be more prevalent in females (59). These results suggest that DNAme differences at these sites may translate to sex biases seen in disease such as Alzheimers and asthma. Moreover, SCAND3 is expressed mainly in seminal vesicle and testis, CRIPS2 is expressed uniquely in testis, whereas GABPA has high expressions in placenta and vagina.

## DISCUSSION

Here, we conducted the first study aiming to characterize autosomal sex differences in DNAme between males and females in whole blood using the IlluminaEPIC BeadChip, which interrogates ~850,000 sites across the genome. Previous studies were performed using Illumina450K BeadChip that covers only ~450,000 sites. We identified 554 sex associated differentially methylated positions on the autosomes whilst adequately handling the technical bias introduced by normalising with the sex chromosomes and correcting for cross hybridisation of some EPIC probes. Previous work has reported contradicting results, with one group finding that there is higher methylation on autosomes in females (5, 21, 60, 61), another group identifying higher methylation on autosomes in males (62, 63) and a third group finding no significant difference in DNAme on autosomes between males and females (19, 64). Our results support the former and we found that 70% of these loci (389 CpGs) showed higher methylation in females compared to males.

The inconsistency seen in the literature is due to the differing normalisation methods applied to DNAme microarray data. Previous research has shown that as the methylation levels of CpG sites on the X chromosome differ largely between males and females, normalisation methods which normalise array data indiscriminately with CpG sites on the autosomes introduce large technical biases for autosomal CpGs (31). Using such normalisation methods, will therefore lead to many autosomal CpG sites being falsely associated with sex. Our choice of normalisation method greatly reduced technical bias at autosomal CpGs for male and female samples, and, thus, we report a highly robust catalogue of autosomal CpGs differentially methylated between males and females.

We further categorized these 554 saDMPs into two groups, those that were hypermethylated in males (n=166) and those that were hypermethylated in females (n=389). Several saDMPs found to be hypermethylated in females overlapped the transcription start site (TSS) of genes not previously been reported to exhibit sex differences in DNAme including DYRK2, SMAD2 and SHANK2. Interestingly, it has previously been shown that sex hormones can regulate SHANK expression leading to a sex differential expression in SHANK2 (65). Furthermore, this gene has previously been implicated in autism spectrum disorder, a disorder known to exhibit higher prevalence in males rather than females (66) In contrast, the most significant saDMP hypermethylated in males is located in the CpG island of a gene located on chromosome 21 called GABPA. GABP is a methylation sensitive transcription factor and has previously been shown to be a transcriptional activator of Cyp 2d-9, which is a gene encoding a male specific steroid in mice (67). Sex differences in these regions have previously been identified (21).

Interestingly, as well as DYRK2 and GABPA being the genes annotated to the most significant saDMPs hypermethylated in females and males respectively, they were also the two most significant saDMRs, suggesting these regions could account for important sex biases observed in some diseases. This is further supported by the fact that GABPA has also been heavily associated with early onset of Alzheimers disease, Parkinsons disease, breast cancer and autism (68) and DYRK2 implication in cancer (69). Our KEGG term analysis at those saDMPs hypermethylated in females also implicated many cancer related KEGG terms including thyroid cancer, endometrial cancer, non-small cell lung cancer and more (Figure 2B).

The saDMR harbouring the highest number of CpG sites (n=58) is located on chromosome 6, overlaps SCAND3/ ZBED9 and is hypermethylated in males. Hypermethylation of this gene has been suggested to be implicated in hepatocellular carcinoma which is third leading cause of deaths related to cancer (70). These results collectively support the hypothesis that sex differences in autosomal DNAme may account for some of the sex differences seen in disease prevalence, onset and progression. Moreover, we did identify saDMPs in genes known to exhibit sex differences in DNAme such as CRIPS2, SLC9A2, DDX43 which are involved in spermatogenesis and male fertility (21, 30). Specifically, CRIPS2 harboured 9 significant saDMPs, all hypermethylated in females, and is part of a group of proteins called CRISPs which show male biased expression in the male reproductive tract. CRIPS2 plays an important role in spermatogenesis, acrosome reaction and gamete fusion (71). Some of our saDMPs were located in genes known to show sex by age effects, such as PRR4, a gene associated with dry eye syndrome (Perumal et al,. 2016). Despite this, recent research shows that the adult blood DNA methylome is largely affected by sex, but that these methylome sex differences do not change throughout adulthood and so are largely independent from age effects (72, 73).

The Illumina EPIC array has an increased coverage of the genome, including distal regulatory elements (74). It was interesting that the 554 saDMPs were still found to be significantly enriched at CpG islands and CpG shores but depleted in open sea regions of the genome (Figure 1E). As the genomic location of DNA methylation normally alters its function, with methylation in CpG islands normally functioning to serve long term silencing of genes (75) and CpG island shore methylation being strongly related to gene expression (76) this suggests a potential functional role for these saDMPs. To further support these findings, we identified enrichment of these saDMPs at enhancers, 5’UTRs and promoters (Figure 1F). Despite this enrichment at regulatory regions, we found little correlation of these sites alone with differences in gene expression between males and females, suggesting that these saDMPs may not be sufficient alone to predict gene expression. It is worthwhile mentioning that, while some genes associated with these saDMPs displayed statistically significant difference in expression between males and females, the differences in expression were modest.

Instead, these saDMPs are likely to interact with transcription factors and other regulatory features to affect gene expression and chromatin organisation. This potential link was identified in our TF motif analysis, where we found SRY (sex determining region Y) and SOX9 (SRY-box 9) transcription factor motifs also known as the sex determining factors, to be enriched at those saDMPs hypermethylated in females and further identified these as hubs in the TF network. SRY has been found to bind and repress WNT activation of ovarian genes, and both SRY and SOX9 transcription factors have been shown to bind the promoter regions of many targets of involved in differentiation of the testis (53). Furthermore, we also found SR1 TFBS enriched in the saDMPs hypermethylated in females, a gene known to interact with SOX9 to increase its own expression to propel differentiation of testis beyond SRY activity (53)

It has previously been reported that 3D genome organisation can impact sex biased gene expression through direct and indirect effects of cohesion and CTCF looping on enhancer interactions with sex biased genes (77), Recently, it was shown that with rising oestrogen levels, the female brain exhibits sex hormone driven plasticity and that chromatin changes underlie this (78). Interestingly, by annotating our saDMPs to distal genes using chromatin loops, we were able to identify contacts between saDMPs and three genes HIST1H3A, HIST1H4A and HIST1H4B which are core components of nucleosome, thereby responsible for playing a role in chromatin organisation. We further were able to identify interactions between these three genes and a gene harbouring a saDMP found to be hypermethylated in males called CDYL. These results suggest that although we found DNAme to not be predictive of sex differences in gene expression (Figure S5), these saDMPs may interact with other genes, transcription factors and other epigenetic modifications to direct chromatin organisation and regulatory networks.

Lastly, we acknowledge limited overlap with previous studies yet conclude that this is due to our extremely large sample size (n=1171) and improved handling of sex bias introduced by normalising such data with the sex chromosomes. Both of these factors contribute to our ability to detect true positives and obtain a more robust catalogue of true sex associated autosomal CpGs.

## AVAILABILITY

The code to perform this analysis is available on GitHub https://github.com/livygrant97/ASD_DNAme

## Competing interests

## ACKNOWLEDGEMENT

The authors acknowledge the use of the High Performance Computing Facility at the University of Essex and would like to thank Stuart Newman for his support.

## FUNDING

OAG is funded by the University of Essex. M.K. was supported by the University of Essex and ESRC (grant RES-596-28-0001). L.S. was supported by Medical Research Council grant K013807. N.R.Z. was supported by the Queen Mary University of London.

Measurement of DNA methylation in Understanding Society: The UK Household Longitudinal Study was funded through an enhancement to Economic and Social Research Council (ESRC) grant ES/N00812X/1.

The analysis was facilitated by access to the Ceres high-performance computing cluster at the University of Essex.

## CONFLICT OF INTEREST

The authors declare that they have no competing interests.

---

**Figure S1:**
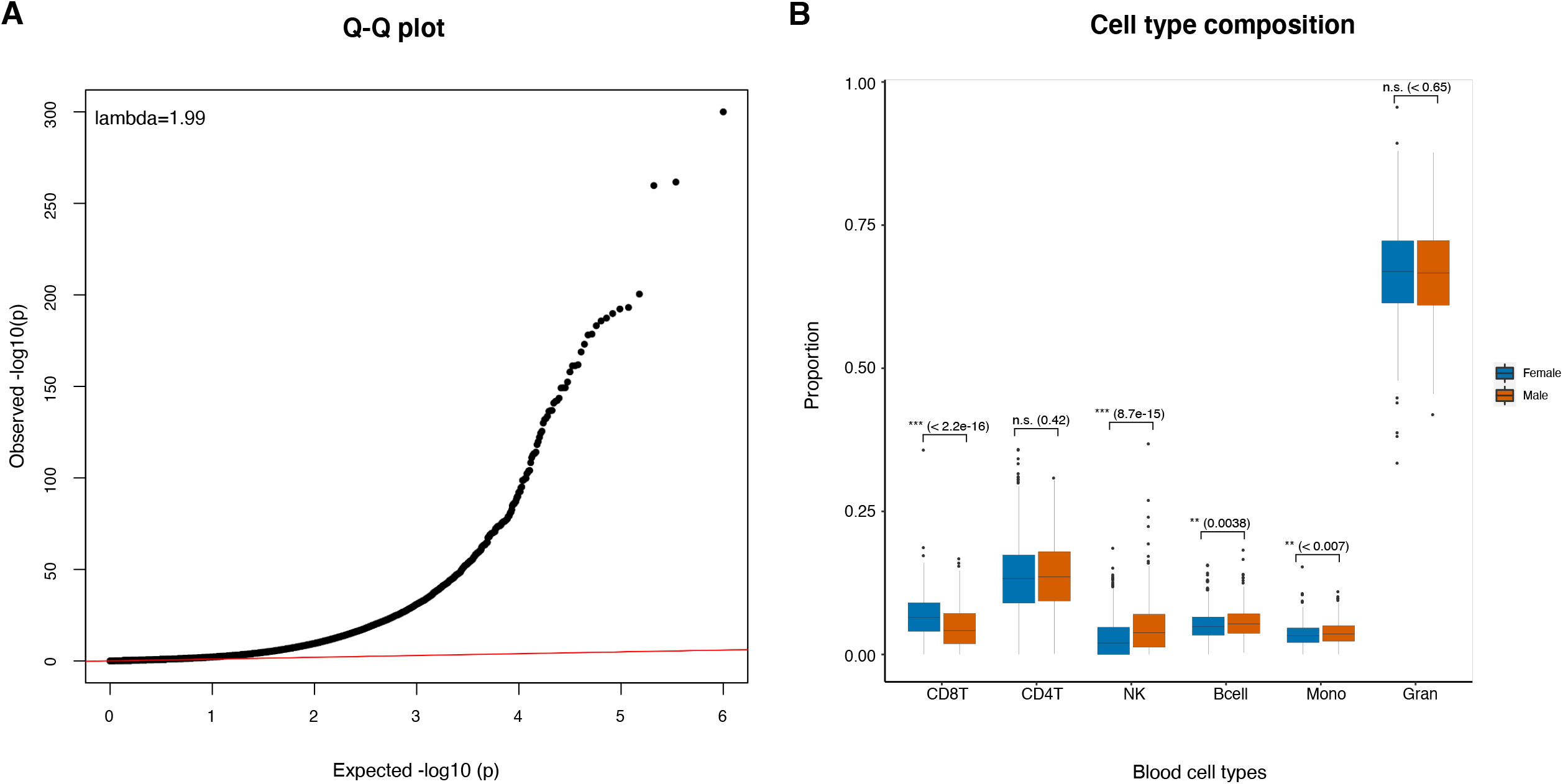
(A) QQ plot and lambda values for the distribution of the adjusted p values against the null distribution for EWAS of sex in the understanding society cohort. Genomic inflation lambda score is indicated in the QQ plot to indicate statistical inflation of p-values. (B) Boxplots of estimated whole blood cell type proportions for males (orange) and females (blue), estimated using the *estimateCellCounts* function from bigmelon. We performed a Mann-Whitney U test (p-value: n.s. ≥ 0.05, * p-value < 0.05, ** < 0.01 and *** < 0.001).

**Figure S2:**
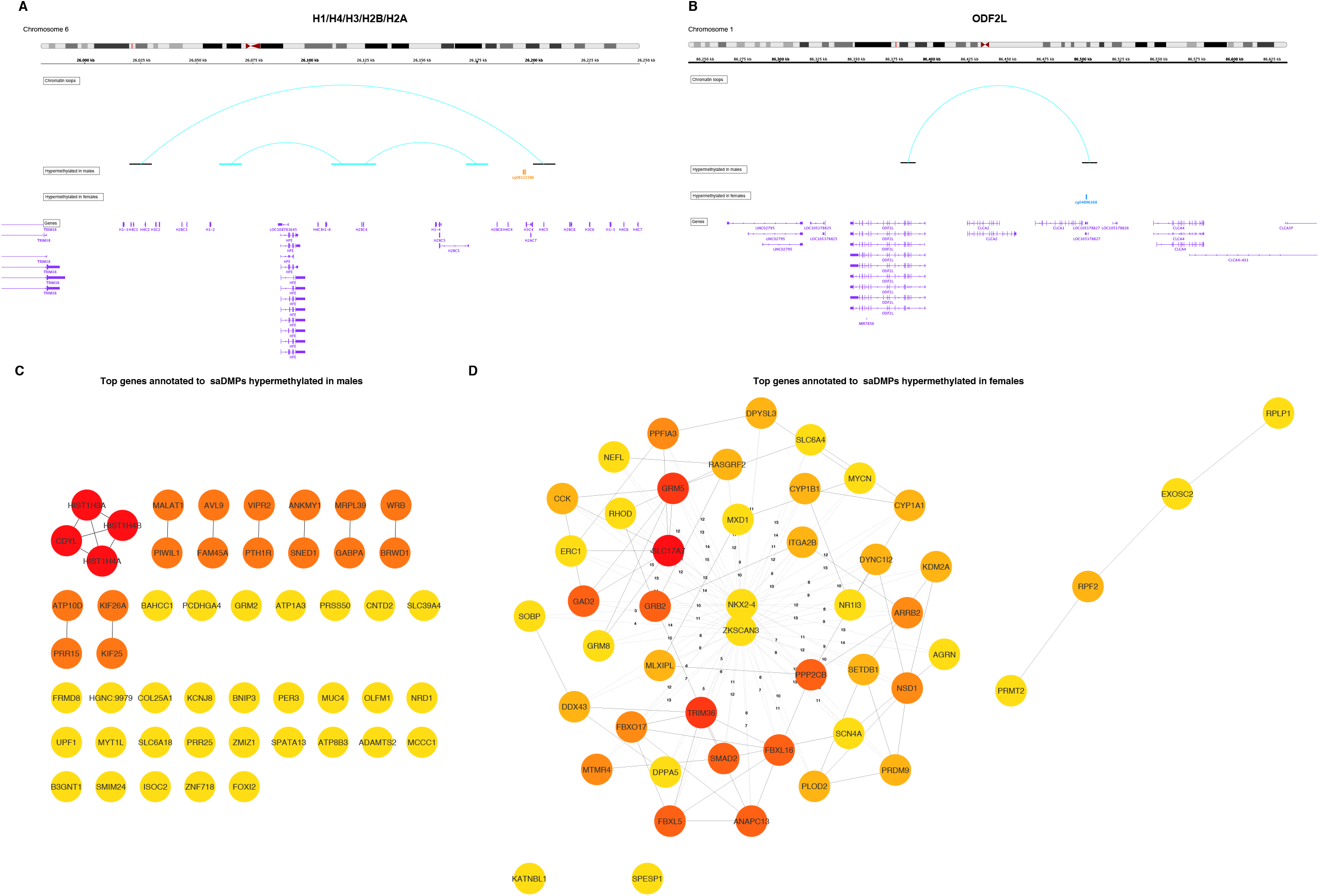
(A) Integrated genomics viewer track of chromatin loop on chromosome 6 showing two saDMPs hypermethylated in males contacting H1/H4/H3/H2V/H2A. (B) Integrated genomics viewer track of chromatin loop on chromosome 1 showing an saDMP hypermethylated in females contacting the ODF2L gene. Blue lines represent the chromatin loops, with black lines showing the loop anchors. Orange vertical lines represent the saDMPs hypermethylated in males and blue vertical lines represent the saDMPs hypermethylated in females. Purple annotations represent genes. (C-D) Subnetworks of the top 50 genes annotated to saDMPs hypermethylated in males (C) and females (D). Node colour represents the degree of connectivity. The scale from red to yellow represents the top 50 enriched TF motif rank from 1-50, with red indicating highest degree and yellow indicating lowest degree.

**Figure S3:**
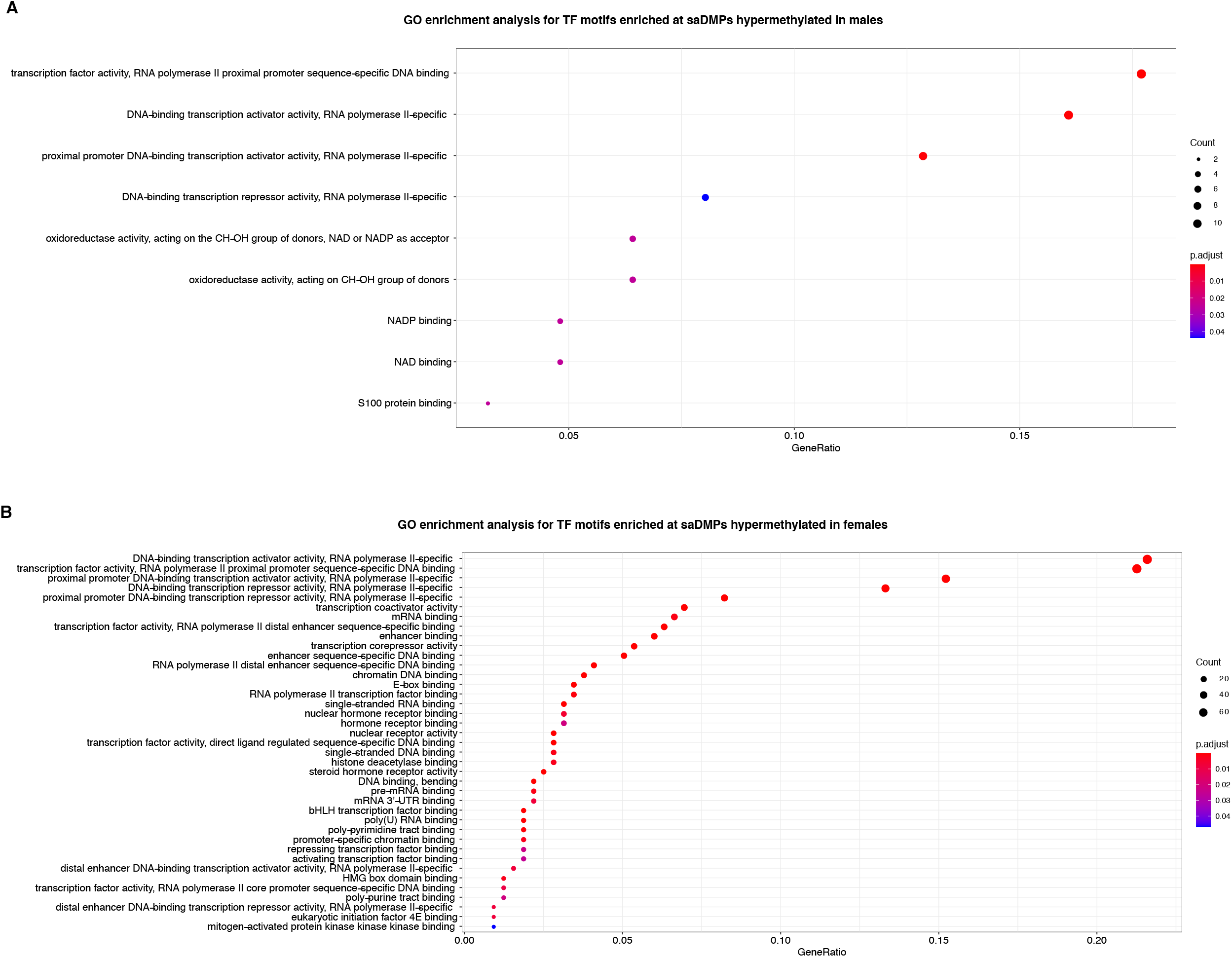
GO terms overrepresented for the significantly enriched TF motifs at saDMPs hypermethylated in males (A) and females (B).

**Figure S4:**
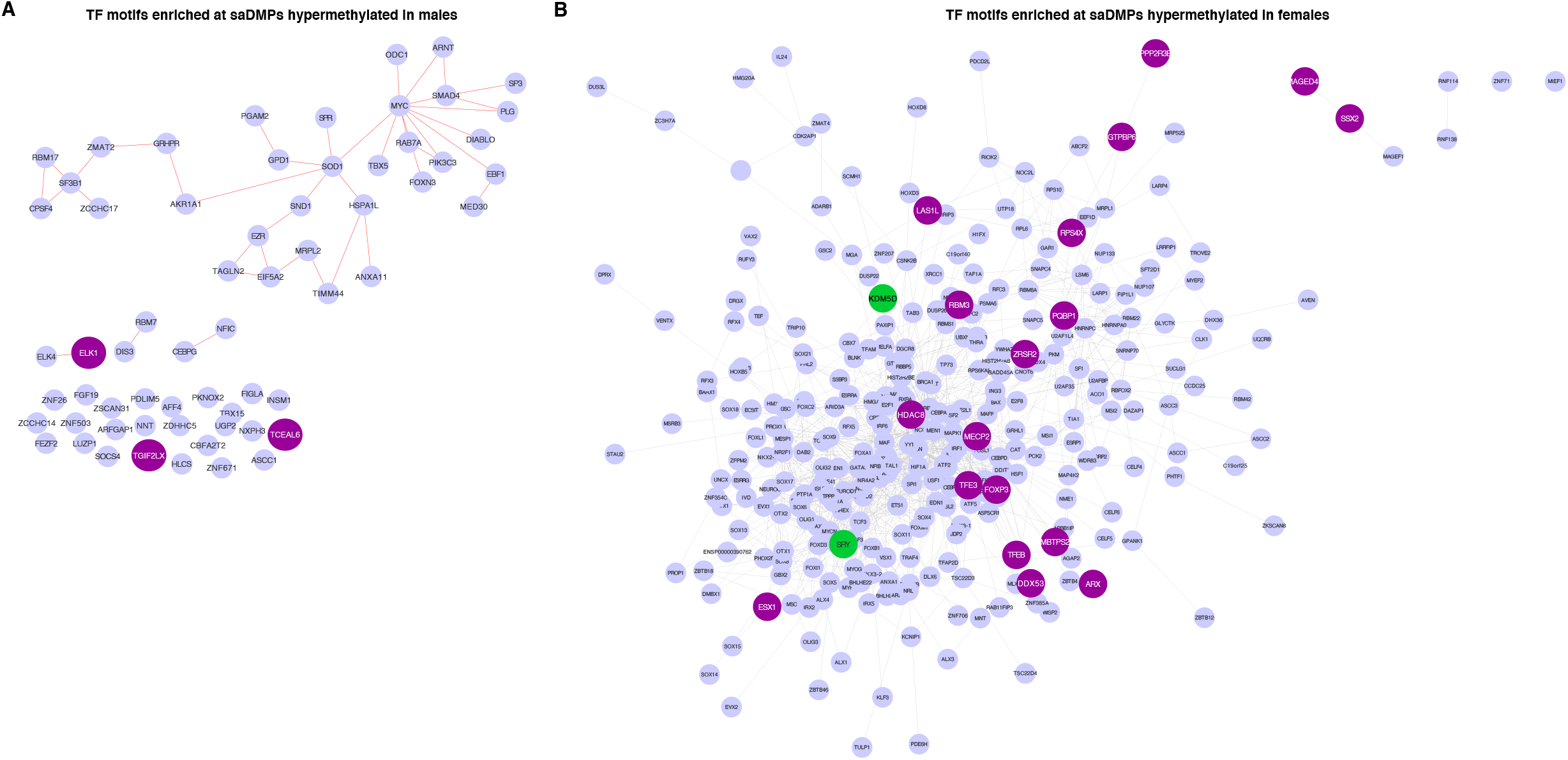
(A-B) Network visualisation of protein-protein interactions for all TF motifs enriched at saDMPs hypermethylated in males (A) and females (B). Grey circles represent individual TFs located on autosomes, while purple circles represent TFs encoded on the X chromosome.

**Figure S5:**
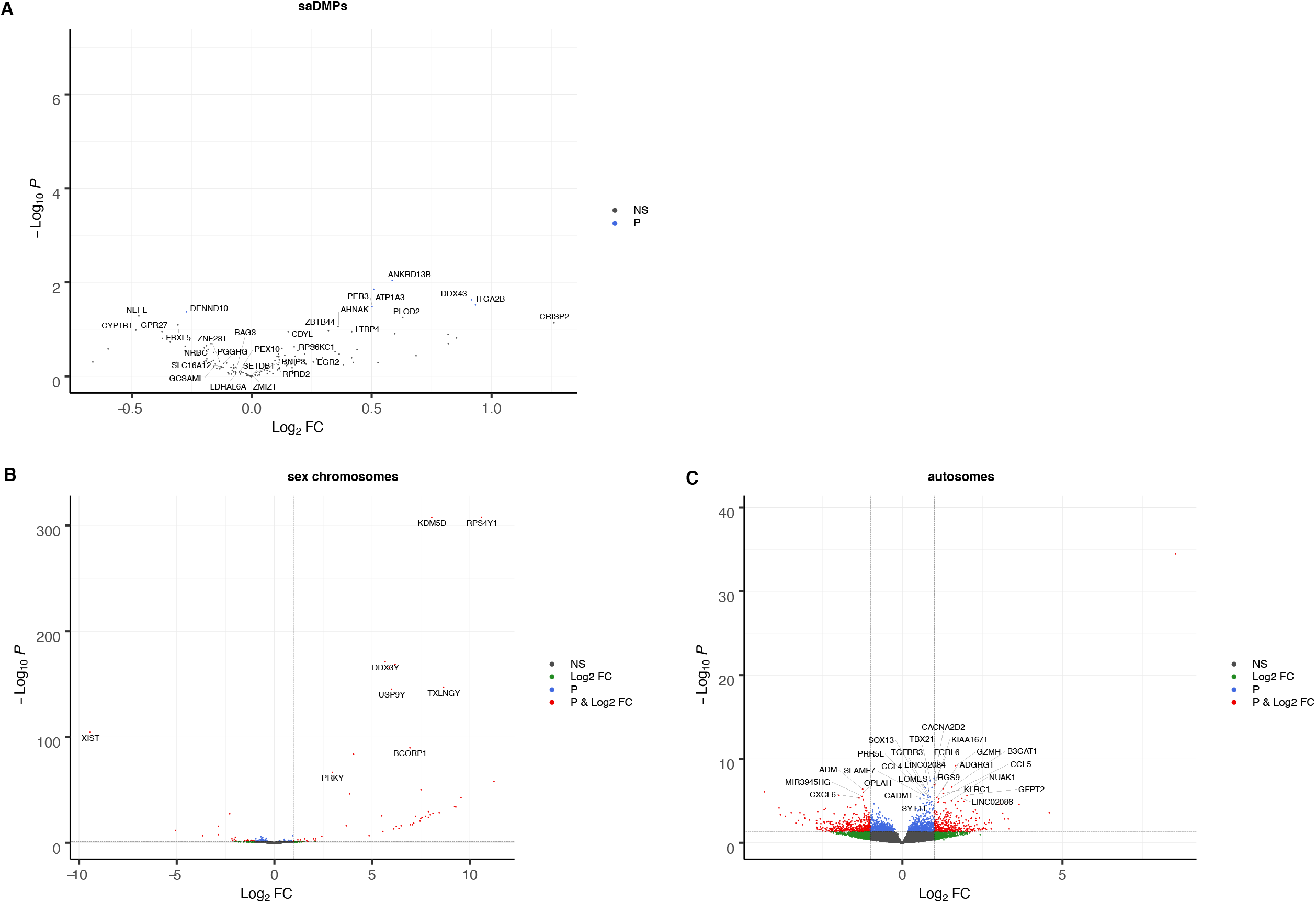
Volcano plot showing differential gene expression between males and females. We considered the case of: (A) genes annotated to the saDMPs, (B) sex chromosome linked genes and (C) autosomal genes. Points coloured in grey represent non differentially expressed genes. Green points represent genes which had a log_2_ Fold Change value greater than 1. Blue points represent genes which met the p value threshold (<0.05). Points coloured in red represent genes which showed differential expression between males and females (p value<0.05 & log_2_FC > 1).

## ADDITIONAL FILES

**Additional file 1: Significant sex associated autosomal DMPs.** Illumina Manifest annotations for all 554 significant CpG sites associated with sex on the autosomes.

**Additional file 2: Significant sex associated autosomal DMRs.** Test results for all significant DMR’s associated with sex on the autosomes ordered by FDR value.

**Additional file 3:** Enrichment statistics for the TF motifs enriched at saDMPs hypermethylated in females.

**Additional file 4:** Enrichment statistics for the TF motifs enriched at saDMPs hypermethylated in males.

**Additional file 5:** Functional enrichment for subnetworks of TF motifs enriched at saDMPs hypermethylated in females.

**Additional file 6:** Functional enrichment for subnetworks of TF motifs enriched at saDMPs hypermethylated in males.

## Notes

### Competing Interest Statement

The authors have declared no competing interest.

